# Modelling mouse auditory response dynamics along a continuum of consciousness using a deep recurrent neural network

**DOI:** 10.1101/2022.04.29.490019

**Authors:** Jamie A. O’Reilly

## Abstract

**Objective:** Understanding neurophysiological changes that accompany transitions between anaesthetized and conscious states is a key objective of anesthesiology and consciousness science. This study aimed to characterize the dynamics of auditory-evoked potential morphology in mice along a continuum of consciousness.

**Approach:** Epidural field potentials were recorded from above the primary auditory cortices of two groups of laboratory mice: urethane-anaesthetized (A, n = 14) and conscious (C, n = 17). Both groups received auditory stimulation in the form of a repeated pure-tone stimulus, before and after receiving 10 mg/kg i.p. ketamine (AK and CK). Evoked responses were then ordered by ascending sample entropy into AK, A, CK, and C, considered to reflect physiological correlates of awareness. These data were used to train a recurrent neural network (RNN) with an input parameter encoding state. Model outputs were compared with grand-average event-related potential (ERP) waveforms. Subsequently, the state parameter was varied to simulate changes in the ERP that occur during transitions between states, and relationships with dominant peak amplitudes were quantified.

**Main results:** The RNN synthesized output waveforms that were in close agreement with grand-average ERPs for each group (r^2^ > 0.9, p < 0.0001). Varying the input state parameter generated model outputs reflecting changes in ERP morphology predicted to occur between states. Positive peak amplitudes within 25 to 50 ms, and negative peak amplitudes within 50 to 75 ms post-stimulus-onset, were found to display a sigmoidal characteristic during the transition from anaesthetized to conscious states. In contrast, negative peak amplitudes within 0 to 25 ms displayed greater linearity.

**Significance:** This study demonstrates a method for modelling changes in ERP morphology that accompany transitions between states of consciousness using a RNN. In future studies, this approach may be applied to human data to support the clinical use of ERPs to predict transition to consciousness.

## 1. Introduction

There is an extensive history of clinical research using noninvasive evoked electrophysiology measures in efforts to understand the extent of neurological function in patients who may be unresponsive due to trauma or administration of dissociative anesthetics [1]–[6]. A general pattern of diminished cortical responsiveness to external stimuli characterizes the transition from consciousness to deepening anesthesia, as seen in auditory event-related potential (ERP) amplitudes [7], [8]. However, some studies suggest that specific components of the ERP are more effective than others for these purposes, such as those in the middle-latency range [9], [10], or those reflecting auditory novelty responses [8], [11], which are found to be differentially affected by state of awareness. A commercial system has been produced that relies on calculating the ratio of amplitudes in small segments of the middle-latency auditory-evoked potential (MLAEP), with lower values indicating no brain activity and higher values indicating greater wakefulness [12]. In relation to auditory novelty responses, the mismatch negativity (MMN) and involuntary P300 (P3a) components are thought to index return to consciousness in unresponsive patients [11].

Given inherent difficulties in ascertaining the underlying neurophysiology reflected by these noninvasive measurements in humans, this research benefits from using experimental animals [13]– [23]. Preclinical studies accommodate more frequent use of invasive recording technologies and greater degrees of manipulation than are typically possible with human subjects, which ought to help decipher precisely how specific component(s) are influenced by genetic, pharmacological, environmental, and behavioral conditions. However, this raises the formidable task of linking components of ERP waveforms extracted from different species; a subject that is being actively pursued [24]. Perhaps there is sufficient rationale for relating ERP components recorded from humans with some non-human primates, although the majority of studies in this field are conducted using rodents, which, for anatomical and physiological reasons, have auditory responses that are less compatible. Despite these challenges, auditory neurophysiology research in rodents continues to supply insights that are of value to human health science. As such, the present article describes an approach in mice that has relevance to the examination of ERP morphology changes in different states of awareness, which potentially has future clinical applications.

Many auditory neurophysiology studies in rodents employ urethane anesthesia because it maintains relatively constant cardiorespiratory function over extended periods [13], [14], [25]–[28]. Auditory-evoked potentials measured from urethane-anaesthetized rodents are typically dominated by a large early stimulus-onset component [14], [25], [28], underpinned by sparse laminar activity [27]. There is also evidence that urethane-anesthetized mice exhibit long-latency auditory novelty responses [29]. The pathways by which urethane exerts its anesthetic effect involve multiple neurotransmitter systems in a complex, imprecise manner [30], which perturbs normal sensory processing [31]–[33]. Nevertheless, it remains a popular veterinary anesthetic for neurophysiology research. Some studies have compared the influence of different anesthetics on rodent ERPs [13], [19]. In a comparison between ketamine-xylazine and urethane-xylazine anesthesia in two groups of mice, Lee and Jones [13] concluded that urethane preserved greater physiological functioning. In another comparative study, Castoldi et al. [19] found that sevoflurane anesthesia attenuated visual evoked potentials more than ketamine-xylazine anesthesia in two groups of rats. These findings suggest that anesthetic agents can be placed on a spectrum in terms of their effects on sensory neurophysiology.

Ketamine and other NMDA receptor antagonists are delivered in sub-anesthetic doses to study their effects on auditory novelty responses in humans and animals [15], [20], [34]–[38]. In some cases, these have been combined with urethane to produce dose-dependent attenuation of proposed rodent analogues of MMN [38], [39], agreeing with findings in humans [34] and non-human primates [36]. This indicates that NMDA receptor disruptors, superimposed upon the pharmacological influence of urethane, produce notable changes in sensory neurophysiology. In conscious mice, ketamine has been found to diminish auditory ERP amplitudes at approximately 40 ms and from 50 to 75 ms post-stimulus [15], [20]. In another study, Nakamura et al. [40] recorded auditory novelty responses of anaesthetized and conscious rats, finding that the former displayed lower amplitude, less dynamic ERP waveforms, lacking deviance-related components between 50 and 75 ms that were present in conscious animals. Collectively, these studies paint a somewhat vague picture suggesting that auditory novelty responses in mice are at least partially dependent on intact NMDA receptor function. These characterizations may be made more explicit with computational modelling that can capture changes that occur in the ERP between states induced by different compounds.

The aim of the present study was to apply a hierarchical recurrent neural network (RNN) to model cortical auditory-evoked potentials from urethane-anaesthetized and conscious mice before and after administering ketamine. After fitting this RNN, it was used to simulate changes in ERP morphology that coincide with transitions between states, thereby providing a mechanism for exploring the neurophysiological consequences of individual and compounded anesthetic agents on the auditory-evoked response in mice.

## 2. Methods

### 2.1 Animals

Two cohorts of laboratory mice were used in this study, as described previously [41]. One group of 14 mice (9 male and 5 female) was anaesthetized with urethane (1.5 g/kg i.p.), placed on an isothermal heat pad, and remained unconscious throughout the experiment. Sustained loss of consciousness was confirmed by absence of corneal and pedal reflexes. Animals in this group were aged 14 to 17 weeks (mean = 15.4), and weighed between 20.6 and 32.4 g (mean = 26.1). The other group of 20 mice (9 male and 11 female) was conscious during experiments. These were aged from 29 to 37 weeks (mean = 32.4) and weighted 23.8 to 36 g (mean = 29). Both groups fall within the young to middle-aged adult range for lab mice, which are considered to have normal hearing [42]. Previous articles have reported the surgical procedures performed to implant epidural field potential electrodes to the conscious [43] and anaesthetized groups [44]. All of these procedures were approved by the Animal Welfare and Ethical Review Body, University of Strathclyde, and performed in accordance with the United Kingdom Animals (Scientific Procedures) Act 1986.

### 2.2 Experiment

Both groups of mice were part of a larger study investigating auditory neurophysiology [45]. They were placed inside an electrically and acoustically isolated chamber for recording epidural field potentials while sequences of auditory stimuli were sounded. Anaesthetized mice were positioned facing a single loudspeaker calibrated to the approximate location of their head. Conscious mice were lured into a restraining tube, which was then placed between two calibrated speakers, one facing each end of the tube. Multiple sequences of auditory stimuli were presented before and after administering ketamine (10 mg/kg i.p.). Animals were temporarily removed from the recording chamber to administer ketamine. In the present study, we will evaluate responses to a consecutive repetition sequence, where a 100 ms, 10 kHz, 80 dB pure-tone stimulus was repeated 100 times consecutively with an inter-stimulus interval (ISI) of 450 ms. This was the first sequence played to both groups at the beginning of the experiment and again following ketamine administration. Auditory-evoked responses were grouped into the following four conditions: conscious (C), conscious plus ketamine (CK), anaesthetized (A), and anaesthetized plus ketamine (AK).

### 2.3 Event-related potentials

Data from animals without gross artifacts were retained for analysis, which meant that three animals were dropped from the conscious group, due to contamination by large artifacts. Epidural field potentials were band-pass filtered between 0.1 and 100 Hz, and then down-sampled to 200 Hz. Trials were segmented from 100 ms before to 500 ms after stimulus onset, with the pre-stimulus average subtracted from the whole epoch to perform baseline correction. Signals recorded from left and right auditory cortex channels were not significantly different, and were thus averaged together to enhancesignal-to-noise. This produced 100 segments of 301 time-samples for each mouse. An ERP was computed for each animal by averaging these trials. Grand-average ERPs for each condition (AK, A, CK, and C) are displayed in Figure 2a-b. Time-wise paired t-tests with Benjamini-Hochberg [46] corrections for multiple comparisons were performed to analyze the effects of ketamine on ERP waveforms. Sample entropy [47] was calculated for each ERP with an embedding sequence length m = 1 and tolerance of r = 0.2; these measurements are plotted by condition in Figure 2c.

### 2.4 Modelling

The modelling approach is illustrated in Figure 1. A hierarchical recurrent neural network (RNN) with four 64-unit hidden layers and a single 1-unit output layer was used to model cortical auditory-evoked response data. Hidden units had the rectified linear unit (relu) activation function, and the output unit had linear activation. Input features for the model were constructed to reflect the auditory stimulus and state in each condition. This consisted of a two-element sequence of 301 samples. The first element of the input feature sequence comprised a unit-step pulse function that was high from 0 to 100 ms, designed to represent the auditory stimulus, which was identical in all four conditions. The second element of the input sequence was a constant value representing state: AK = 0.25, A = 0.5, CK = 0.75 and C = 1. These values do not necessarily imply that a linear relationship exists between conditions. Trials from each animal were averaged together to produce an ‘idealized experiment’, with 100 trials in each condition, which were provided as targets for the model. The model was trained to minimize mean-squared error (MSE) between its outputs and these cortical responses. As such, its outputs in response to the four input conditions converged towards the grand-average ERPs, as shown in Figure 3. Adaptive moment estimation (adam) optimization was used with a learning rate of 0.001, β1 of 0.9 and β2 of 0.99. Glorot uniform initialization [48] was used for weights connecting layers, whereas orthogonal initialization [49] was used for recurrent weights. The RNN was trained with backpropagation through time (BPTT) for 1000 steps and batch size of 32, as implemented in Tensorflow [50].

**Figure 1.**
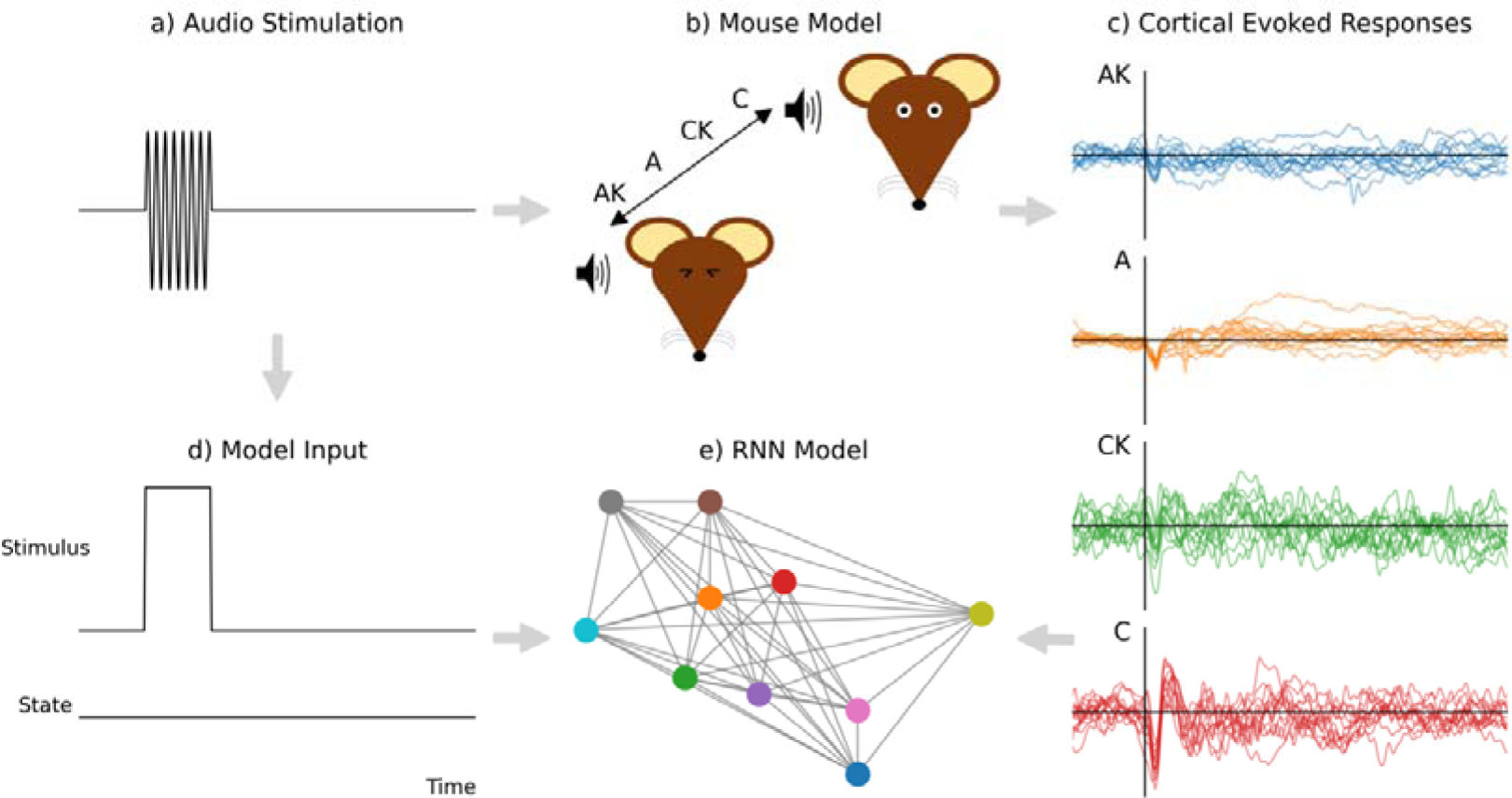
Approach used to fit a computational model to auditory neurophysiology data from anaesthetized and conscious mice. a) An identical sequence of 100 repetitions of a 100 ms, 10 kHz, 80 dB pure-tone stimulus with inter-stimulus interval of 450 ms was delivered. b) Two groups of mice were used: i) anaesthetized (A; n = 14), and ii) conscious (C; n = 17); both were also administered ketamine at 10 mg/kg i.p. (AK and CK), ordered along a ‘continuum of consciousness’. c) Cortical auditory evoked potentials were recorded from mice during audio stimulation. These were subsequently used as labels for training a recurrent neural network (RNN). d) Audio stimuli were represented as a unit step pulse and state was encoded as an unchanging scaler value. e) The RNN model was trained with input features representing sounds and labels from mouse cortical evoked responses.

**Figure 2.**
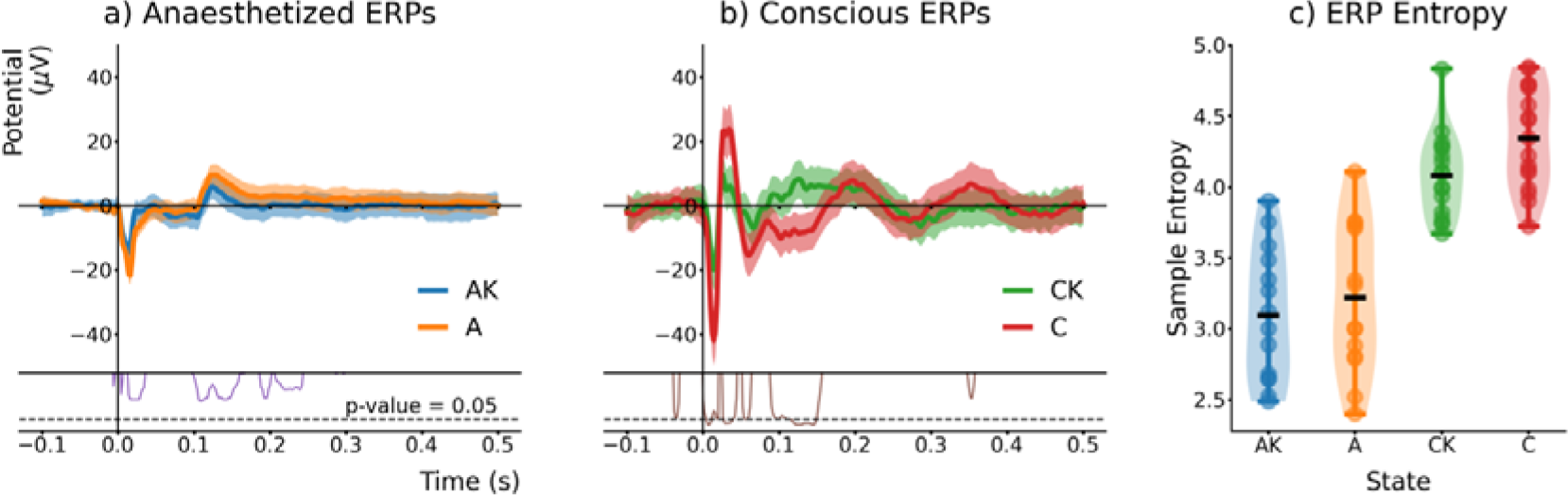
Conscious mice display a more dynamic ERP than anaesthetized mice. a) The response of anaesthetized mice consists of negative stimulus-onset and positive stimulus-offset peaks that are marginally diminished following ketamine administration. b) Conscious mice exhibit larger amplitude, more dynamic auditory responses, resembling a damped oscillation of alternating polarity components, which are significantly attenuated following ketamine. Shaded error bars represent standard error. Lower inset axes in (a) and (b) display corrected p-values from paired t-tests with FDR corrections; the demarcation below which *p* < *α* is indicated with a dashed horizontal line. c) Sample entropy measurements used to arrange responses along a ‘continuum of consciousness’, reflecting a gradient of physiological responsiveness, from anaesthetized plus ketamine (AK), anaesthetized (A), conscious plus ketamine (CK), and conscious (C) mice. Black horizontal bars depict group means.

**Figure 3.**
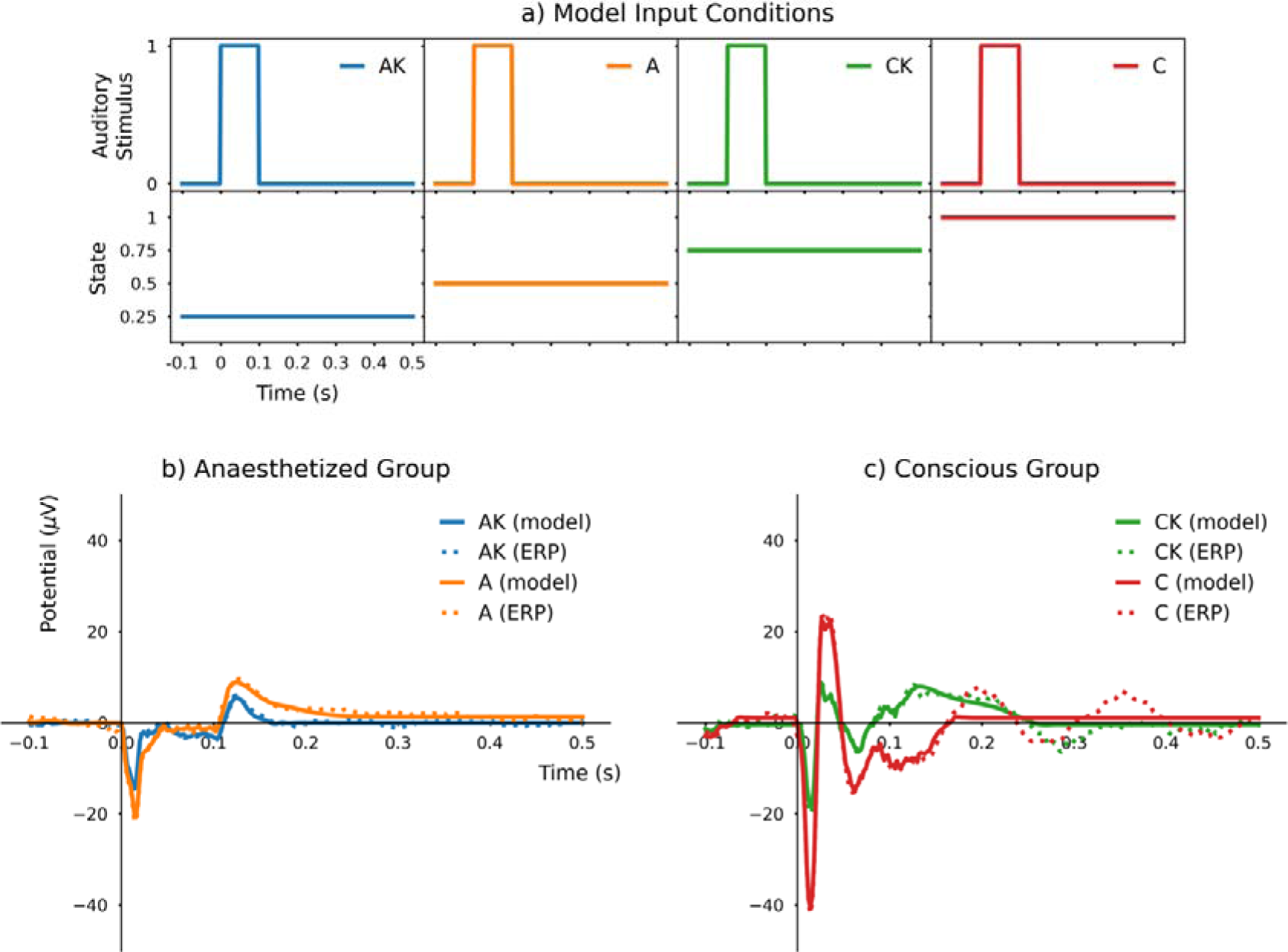
Recurrent neural network accurately models auditory ERP of anaesthetized and conscious mice with and without ketamine. a) Model inputs comprised sequences of two elements, with one representing auditory input and the other representing state. b) Outputs from the model plotted alongside grand-average ERPs from anaesthetized mice before and after ketamine (A and AK, respectively). c) Model outputs and grand-average ERP waveforms from conscious mice before and after receiving ketamine (C and CK, respectively). Beyond approximately 200 ms post-stimulus, amplitude fluctuations in conscious mice ERPs are not captured by the model.

The model architecture was cross-validated over five folds, where 20% of the idealized trials were randomly selected for preparing validation ERPs (one for each condition), and the remaining 80% of trials were used for training data. Models were optimized over 100 iterations, then their outputs were compared with training and validation ERPs using Pearson’s correlation coefficient (r^2^). This procedure yielded average overall performance of r^2^= 0.906 for the training set and r^2^= 0.814 for the validation set (both p < 0.0001), demonstrating a reasonable degree of generalization.

### 2.5 Simulation

The purpose of this simulation was to examine changes in auditory response morphology predicted to occur in the transitions between and beyond states. To achieve this, 101 different input sequences with state values ranging from 0 to 1.25 were synthesized and used to generate outputs from the RNN. In each case, the first element in the input sequence, representing the auditory stimulus, was identical. Resulting simulated ERP waveforms are shown in Figure 4a. Component morphology changes were characterized by measuring peak amplitudes within the measurement windows 0 to 25 ms (N1 peak), 25 to 50 ms (P1 peak), and 50 to 75 ms (N1 peak), which are plotted respectively in Figure 4b-d. A straight line was fitted to N1 peak amplitude measurements, whereas logistic functions were fitted to P1 and N2 peak amplitudes. Parameters for these linear and logistic functions were determined by least squares optimization. These curve fittings were evaluated using Pearson’s correlation coefficient.

**Figure 4.**
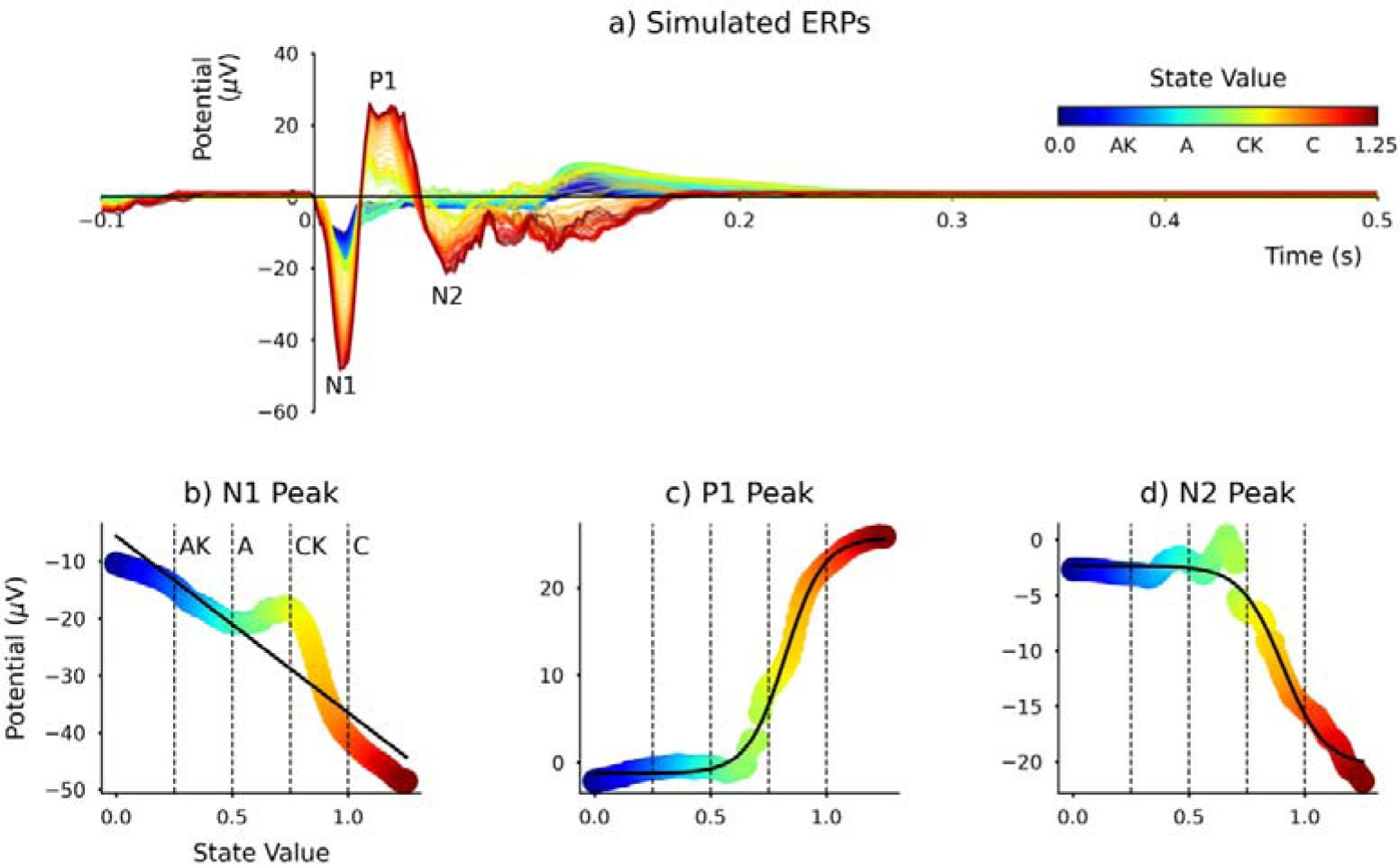
Simulated model outputs portray ERP morphology changes accompanying transitions through the continuum of consciousness. a) Incrementing the model state value input from 0.0 to 1.25, crossing values encoding anaesthetized plus ketamine (AK), anaesthetized (A), conscious plus ketamine (CK), and conscious (C) states, simulates shifting ERP morphologies. b) Negative peak amplitude measured from 0 to 25 ms (N1), demonstrating a persistent negative amplitude across all states. c) Positive peak amplitude measured from 25 to 50 ms (P1), which is absent in anaesthetized state and increases sharply with transition to consciousness. d) Negative peak amplitude measured from 50 to 75 ms (N2), showing a sharp negative increase with transition to consciousness. The solid black line in (b) displays least-squares linear regression, while those in (c) and (d) represent fitted logistic functions. This analysis demonstrates that later components (P1 and N2) tend to extinguish rapidly in the transition from consciousness to anaesthetized states, whereas the earliest component (N1) persists.

### 2.6 Software

Python 3, Matplotlib 3.3.3 [51], MNE 0.23.4 [52], Numpy 1.19.5 [53], SampEn 0.0.17 [47], Scipy 1.7.1 [54], and Tensorflow 2.4.1 [50] were used in this study. The code is available on request.

## 3. Results

Grand-average ERP waveforms shown in Figure 2a-b demonstrate substantial differences in morphology between anaesthetized (A and AK) and conscious (C and CK) groups. The anaesthetized group exhibited prominent negative polarity stimulus-on and positive polarity stimulus-off components, whereas the conscious group displayed a series of alternating polarity peaks, as described previously [41]. Ketamine was not found to produce significant changes to ERPs from urethane-anaesthetized mice, but did significantly diminish component amplitudes from stimulus-onset response through to 150 ms post-stimulus in conscious mice. Sample entropy measurements of ERPs from animals in each condition are shown in Figure 2c. Conscious mice tended to have ERPs with higher entropy than anaesthetized mice. Moreover, ERPs recorded following ketamine administration tended to exhibit less entropy than those preceding ketamine for both groups. Ordered by increasing mean sample entropy of ERP waveforms, conditions AK ⍰ A ⍰ CK ⍰ C may be conceptualized as reflecting a continuum of consciousness. This is not intended to be a rigid or comprehensive taxonomy, nor is not assumed to be linear or reflective of the entire diversity of potential states of awareness that mice are capable of experiencing. Rather, this ordered spectrum of states provides a convenient and rational basis for modelling transitions in ERP morphology observed from mice in each condition of this study.

Model inputs representing the four conditions are illustrated in Figure 3a. Resulting model outputs are plotted alongside grand-average ERP waveforms in Figure 3b-c. Model outputs strongly correlate with the grand-averages, with Pearson’s correlation coefficients calculated for AK (r^2^ = 0.98, p = 2.2 × 10), A (r^2^ = 0.986, p = 3.83 × 10), CK (r^2^ = 0.937, p = 1.06 × 10), and C (r^2^ = 0.937, p = 2.42 × 10). In the conscious group, model outputs deviate from the grand-average ERP beyond approximately 200 ms post-stimulus; subsequent low-amplitude variations of the ERP recorded from conscious mice before administering ketamine are not recapitulated by the model.

The trained RNN was used to simulate ERP waveforms while varying the state parameter of the input sequence. Output waveforms are plotted in Figure 4a. This simulation predicts changes in ERP morphology that occur during the transitions between different states. Peak measurements from prominent features of the simulated waveforms are plotted in Figure 4b-d. The first negative deflection (N1) was fitted with the linear function, *y* = −30.95x – 5.56 (r^2^= 0.923, *p* = 5.79 × 10^−43^), where *y* is the peak amplitude and *x* is the input state value. In contrast, measurements of the first positive deflection (P1) were characterized by a logistic function, *y* = 26.94 / (1 + exp(−12.28(x – 0.82))) – 1.2 (r^2^ = 0.997, *p* = 9.76 × 10–116). The second negative deflection (N2) showed an inverted logistic trajectory, with *y* = −18.04 / (1 + exp(−11.08(x – 0.9))) – 2.33 (r^2^ = 0.987, *p* = 1.04 × 10–80). The straight line fitted to N1 peak amplitude measurements shown in Figure 4b is less strongly correlated than the sigmoidal curves fitted to P1 and N2 peak amplitudes displayed in Figure 4c-d, largely due to the ‘kink’ induced by the CK condition. Nevertheless, it appears evident that the N1 component persists in the transitions between states, whereas P1 and N2 components are almost entirely abolished in the anaesthetized conditions, precipitated by changes following ketamine administration to conscious mice.

## 4. Discussion

The ERP from urethane-anesthetized mice exhibits markedly different morphology from that of conscious mice (Figure 2a-b). The first negative peak (N1: 0 to 25 ms) is apparent in both groups, and then conscious mice display a series of alternating polarity peaks (P1: 25 to 50 ms, and N2: 50 to 75 ms). In both cases, ERP amplitudes were diminished following ketamine, although only the conscious group was found to exhibit statistically significant differences across the N1, P1 and N2 latency ranges, and a subsequent window from 100 to 150 ms post-stimulus. Polarity inversion of these three peaks relative to previous studies [20], [40] is considered to arise from differences in electrode configuration. Despite displaying opposite polarities, the latencies of these three prominent peaks agree with prior findings in rodents. It is noteworthy that stimuli in this experiment were presented in a consecutive repetition sequence, thereby presumably not evoking auditory novelty components that are expected from oddball stimulation. This may be taken to suggest that ketamine ablates obligatory components of the auditory ERP in mice.

Sample entropy (Figure 2c) was used to evaluate conditions according to the average information content present in their ERP signals [47]. By doing so, this provided supporting rationale for ordering conditions that followed perceived levels of awareness; i.e. in ascending order, AK ⍰ A ⍰ CK⍰ C. Encodings for each state were derived based on this order and paired with simulated auditory input for training the RNN (Figure 3a). It is worth stating that this approach does assume a linear relationship among these different states, which would not be justified, given that the precise pharmacodynamics of and interactions between ketamine and urethane are ambiguous. Nevertheless, it is understood that combining multiple anaesthetic substances tends to compound their depressive actions on physiological systems. Also, while the encodings depict a linear distance between states, the RNN is capable of learning nonlinear functions of its inputs. It is evident from the synthetic ERP waveforms generated by the model (Figure 3b-c) that these are in close agreement (r^2^ > 0.9) with grand-average ERPs from mice in each condition. It is feasible that other anaesthetics could produce different ERP morphology changes [13], [19], [55]. Furthermore, more subtle differences in state of conscious awareness (e.g. high-stress vs. low-stress) influences perception, and may therefore differentially affect auditory neurophysiology [56].

At present, we are unaware of this computational modelling approach being applied to characterize ERP waveforms. The majority of applications of deep learning techniques in neuroscience research fall under the category of signal classification or decoding models [57], which can be particularly effective for brain-computer interfacing [58]. Recurrent neural networks trained by BPTT are known to biologically implausible, but nevertheless offer a potentially valuable tool for studying the computational principles underlying cognitive neuroscience and physiology data [59], [60]. In the present study, after modelling the grand-average ERP waveforms, the state encoding input was manipulated to explore the transitions in ERP morphology between states using the trained RNN (Figure 4). In this respect, the RNN was used as a generative model to produce synthetic ERP waveforms. A similar approach may be taken to characterise the influence of anaesthetics on human auditory-evoked responses [7], [8]. Moreover, recent efforts to explore differences in auditory-evoked responses across primates [24] could be supplemented by applying this methodology, if sufficient control of recording conditions can ensure a high degree of comparability.

Ketamine and other NMDA receptor antagonists have long been known to diminish auditory novelty response amplitudes in rodents [20], [37]–[39], non-human primates [36], and humans [34]. Results of the present study add to this literature by demonstrating that obligatory components of the mouse auditory ERP are also significantly diminished by NMDA receptor disruption. The effects of urethane anesthesia observed on mouse ERPs here are also consistent with general observations of the influence of anesthetics on evoked responses in humans and other animals [7], [8], [13], [19], [55]. Simulated ERPs were generated using state encodings spanning beyond the range of values used during training, allowing a characterization of the dynamics in ERP morphology throughout this continuum of consciousness. The logistic character of P1 and N2 peak amplitude dynamics (Figure 4c-d) indicate that these components are rapidly extinguished with loss of consciousness, whereas the more linear property of N1 peak amplitude dynamics suggest that it is more reliably evoked from unconsciousness animals.

These findings can be interpreted within the context of the predictive coding theory of sensory disconnection postulated by Sanders et al. [61]. As the first and most persistent feature of the observed auditory-evoked response, the N1 peak is considered to reflect a feed-forward process of excitation from the thalamus to primary auditory cortices. Changes in later deflections that were initially present in conscious animals (P1 and N2) may signify loss of connectedness and subsequently consciousness that accompany diminished feed-back sensory predictions from higher-order cortical areas [61]. Another perspective from which to view these results is through the lens of integrated information theory (IIT) [62], [63]. Increasing sample entropy, a proxy for information content in physiological signals [47], measured from conditions spanning the continuum of consciousness, are entirely consistent with the principal notion of IIT. Within this framework, changes in N1, P1 and N2 observed from the simulation may be interpreted to reflect hypothetical changes in Φ (phi), the integrated information correlate of consciousness.

One perceived limitation of this study may be that two separate groups of mice were used, and these were of slightly different age ranges. The conscious group were older than the anesthetized group, although both were considered to be young to middle-aged for laboratory mice, with no suspected hearing loss at the intensity level of stimuli used in this study [42]. Moreover, ERPs from the conscious group had larger, more dynamic amplitude changes than those of the anesthetized group. This is precisely the opposite of what would be expected if the differences between groups were principally due to age-related hearing loss [42]. Aside from this, other studies have also considered using different cohorts of animals in treatment groups to be a valid approach [13], [19]. As such, while it would have been preferable to use the same animals in all four conditions, this caveat does not demand a substantial reinterpretation of the results. Perhaps a more consequential caveat is that other sequences of sound stimulation were delivered to animals between the initial recording and subsequent recording after administering ketamine. As such, we cannot completely extricate changes in ERP morphology between pre-and post-ketamine responses that might be due to neurophysiological adaptation, which is known to unfold across multiple timescales [64], [65].

## 5. Conclusion

This study has demonstrated a computational method for simulating the dynamics of ERP morphology changes that occur in the transitions between states of awareness placed along a hypothetical ‘continuum of consciousness’. Being applied to cortical auditory evoked potential data from urethane-anesthetized and conscious mice, before and after administering ketamine (10 mg/kg i.p.), this modelling approach depicts relative preservation of N1 (0 to 25 ms) peak amplitude, and comparatively diminished P1 (25 to 50 ms) and N2 (50 to 75 ms) amplitudes, in the transition from consciousness towards unconsciousness, mediated by ketamine and then urethane. It would be feasible to apply this method to model data from patients, which could find fruitful clinical application in characterizing noninvasive electrophysiological biomarkers in anesthesiology and consciousness science.

## Acknowledgements

I wish to thank Dr. David M. Thomson of Strathclyde University for training on the preparation and delivery of ketamine to laboratory mice. Thanks also to Dr. Jordan Wehrman of Sydney University for comments on an earlier version of this manuscript. The data used in this study were recorded with support from a doctoral training award from the UK Engineering and Physical Sciences Research Council [grant no. EP/F50036X/1], and the RNN was trained using a graphics processing unit purchased with support from the Research Institute of Rangsit University [grant number 90/2561].

## Data availability

The data and code are available on request.

## Conflict of interest

The authors declare no conflicts of interest.

## Notes

### Competing Interest Statement

The authors have declared no competing interest.

